# Refinement of metabolite detection in cystic fibrosis sputum reveals heme negatively correlates with lung function

**DOI:** 10.1101/475525

**Authors:** Nathaniel R. Glasser, Ryan C. Hunter, Theodore G. Liou, Dianne K. Newman, Mountain West CF Consortium Investigators

**Affiliations:** Division of Biology and Biological Engineering, California Institute of Technology, Pasadena, California; Department of Microbiology and Immunology, University of Minnesota, Minneapolis, Minnesota; The Center for Quantitative Biology and The Adult Cystic Fibrosis Center, Division of Respiratory, Critical Care and Occupational Pulmonary Medicine, Department of Internal Medicine, University of Utah, Salt Lake City, Utah

## Abstract

*Pseudomonas aeruginosa* lung infections are a leading cause of morbidity and mortality in cystic fibrosis (CF) patients (1, 2). Our laboratory has studied a class of small molecules produced by *P. aeruginosa* known as phenazines, including pyocyanin and its biogenic precursor phenazine-1-carboxylic acid (PCA). As phenazines are known virulence factors (3), we and others have explored the possibility of using phenazine concentrations as a marker for disease progression (4–6). Previously, we reported that sputum concentrations of pyocyanin and PCA negatively correlate with lung function in cystic fibrosis patients (6). Our study used high performance liquid chromatography (HPLC) to quantify phenazines by UV–vis absorbance after extraction from lung sputum. Since our initial study, methods for metabolite analysis have advanced considerably, aided in large part by usage of mass spectrometry (LC-MS) and tandem mass spectrometry (LC-MS/MS). Because a more recent study employing LC-MS/MS revealed a surprising decoupling of *P. aeruginosa* metabolites in sputum and the detection of *P. aeruginosa* through culturing or microbiome profiles (4), we decided to check whether we could reproduce our previous findings by analyzing sputum samples from a different patient cohort with a new LC-MS instrument in our laboratory. Our new samples were provided by the Mountain West CF Consortium Sputum Biomarker study (7). In the course of performing our new analyses, comparison of our old HPLC data to our new LC-MS data led us to realize that the peak previously assigned to PCA instead originates from heme, and the peak assigned to pyocyanin originates from an as-yet unknown compound. This correction only affects the measurements of phenazines in sputum, and we are confident in the phenazine measurements from isolated cultures and the 16S rRNA gene sequencing data from that study (6). Here we outline the basis for our correction and present additional data showing that heme concentration negatively correlates with lung function in cystic fibrosis patients.

## Results

### Identification of heme in sputum

Motivated by a recent LC-MS/MS study which reported phenazines in only a fraction of samples with *P. aeruginosa* (4), we sought to determine whether we could repeat our previously reported correlation between phenazines and disease progression (measured by FEV_1_%). We obtained fresh sputum samples from the Mountain West CF Consortium (MWCFC) sputum biomarker study and analyzed 71 sputum supernatants using HPLC coupled to UV–vis detection and a single quadrupole mass spectrometer (Waters Acquity QDa). Patients with confirmed CF enrolled in the MWCFC study were randomly selected from all sputum producers followed by the participating centers and enrolled during clinical stability. Patients had a mean age of 28 years (SD=12), FEV_1_% of 70% (SD = 22), weight-for-age *z*-score of −0.17 (SD=0.98) at the time of enrollment. Patients had a history of 1.7 pulmonary exacerbations in the year prior to enrollment (SD=1.7), diabetes in 22%, pancreatic insufficiency in 92.1% and 5-year survival predictions (8) ranging from 9.4% to >99.99% (median 94%). *P. aeruginosa* was recovered from 61% of cultures from enrollment and 74% of cultures obtained during the year of enrollment from the MWCFC patients. There were no significant differences between the characteristics of the MWCFC study group and the same characteristics of patients six years of age and able to produce sputum found in the 2014 CF Foundation Patient Registry (7).

To our surprise, and in contrast to our prior study, we were unable to detect the phenazines pyocyanin, PCA, phenazine-1-carboxamide, or phenazine-1-hydroxide in any of the samples to within our detection limit (approximately 0.1 µM) in either the UV–vis or mass channels. Instead, we observed a late-eluting compound in all 71 samples which had an absorbance spectrum that overlaps with the phenazine spectrum but is distinguished by a maximum at 398 nm. This compound produced positive ions at an *m*/*z* of 616.2, further distinguishing it from any known phenazine, and it was present even in patients who did not test positive for *P. aeruginosa*. We noted a striking similarity between the UV–vis spectrum of this compound with published spectra for the porphyrin ring of heme-like compounds (9). We therefore wondered whether this peak represented a compound related to heme, and whether we might have misassigned it previously in sputum samples, which are more chemically complex than bacteriological media.

To further interrogate the peak, we transferred our method to an instrument coupled to a high-resolution quadrupole time-of-flight mass spectrometer (Waters Xevo). Again we consistently observed a UV–vis chromatographic peak at 398 nm (Figure 1A). Consistent with its identity as a heme, this peak had a retention time identical to that of a hemin standard (Figure 1A). Both the sputum samples and the hemin standard gave an equivalent peak in the extracted ion chromatogram for an *m*/*z* of 616.1773 ± 0.01 (Figure 1A), the *m*/*z* expected for hemin having lost its associated chloride ion to produce ferriprotoporphyrin IX. We note that the hemin standard was dissolved in base, a condition known to replace the chloride ion with hydroxide (10) which is subsequently removed in the acidic conditions of our chromatography, therefore explaining the origin of ferriprotoporphyrin IX from our hemin standard. In the positive ion channel (Figure 1B), sputum and hemin produced equivalent signals that were indistinguishable from the expected mass of ferriprotoporphyrin IX to within the resolution of the instrument (∆ppm = 0.97 and −1.6, respectively). Similarly, in the negative channel (Figure 1C), both sputum and hemin produced ions consistent with the formic acid adduct of ferriprotoporphyrin IX (∆ppm = −0.45 and 0.45, respectively), an additive used in our separation method. Application of collision energy produced equivalent fragmentation patterns in the negative ion mode (Figure 1C) with the major daughter ion appearing around 570.1718, consistent with two decarboxylations or a single decarboxylation and loss of formic acid. The sputum peak and the hemin standard also produced indistinguishable UV−vis spectra (Figure 1D) which closely matched previous reports for heme (9). Based on identical retention times, mass and UV–vis spectra, and mass fragmentation patterns, we assign this peak to ferriprotoporphyrin IX, also known as heme B (Figure 1E).

**Figure 1.**
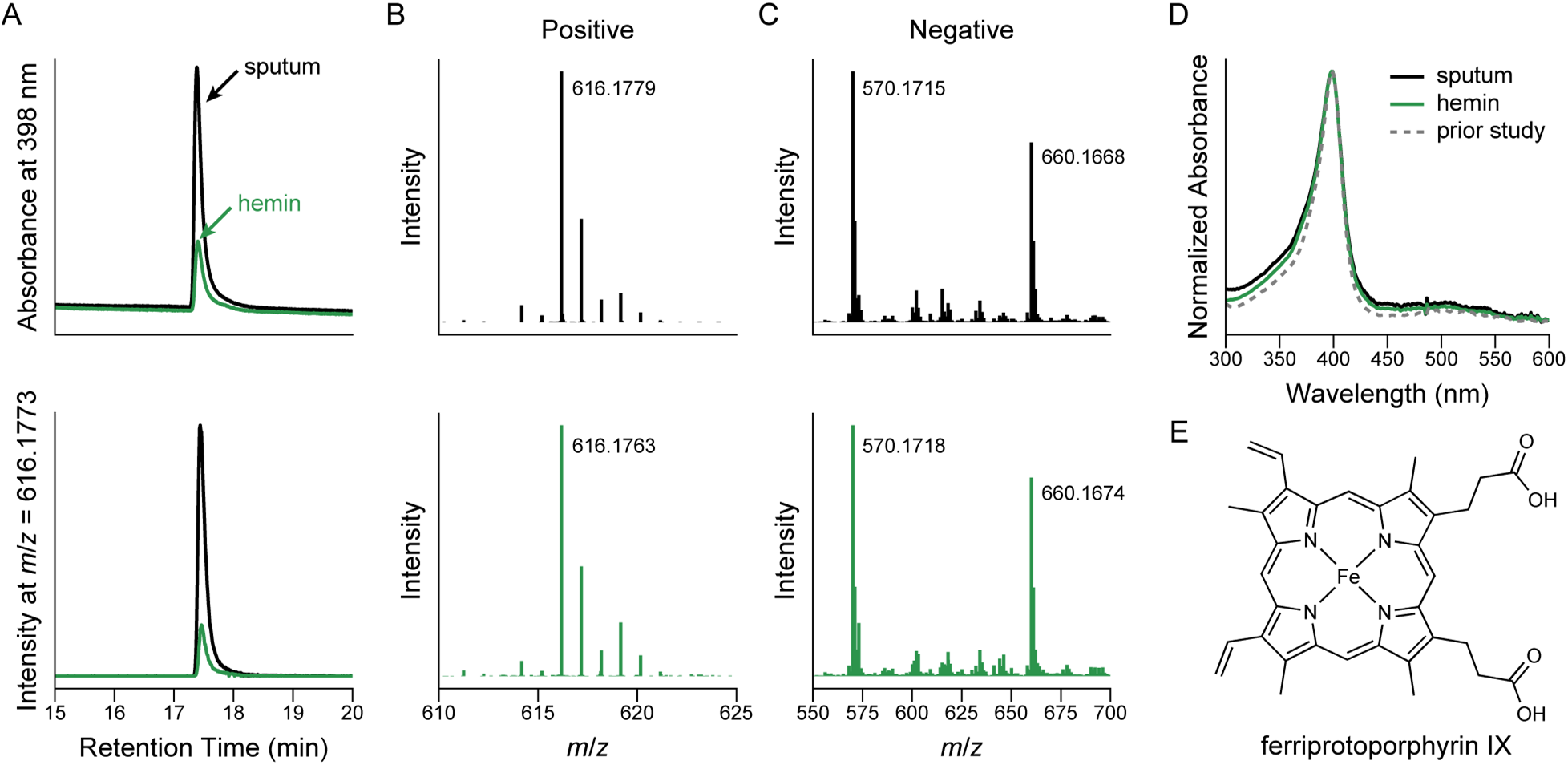
Identification of heme in sputum samples by comparing sputum (black) to a hemin standard (green). (A) UV–vis chromatogram (top) and extracted ion chromatogram (616.1773 ± 0.01 Da) from the positive mass channel (bottom) demonstrating identical retention times. (B) The associated positive ions detected from the peak shown in A, comparing the sputum peak (top) to the hemin standard (bottom). (C) The associated negative ions detected from the peak shown in B, comparing the sputum peak (top) to the hemin standard (bottom). A collision-energy ramp of 10 to 14 eV was applied to generate a fragmentation pattern. (D) Comparison of the UV–vis spectra for the peaks in A (black and green) compared to the same peak identified in our prior study (gray, dashed). For clarity, the spectra are normalized to their maximum value. (E) The chemical structure assigned to the peak in A, ferriprotoporphyrin IX.

### Re-analysis of the original data

Though the MWCFC cohort differed in inclusion criteria from that used in our prior Children’s Hospital Boston (CHB) sputum study (*i.e.* a positive diagnosis of CF based on genotyping or sweat test and positive *P. aeruginosa* culture), we were nevertheless surprised that we were unable to detect phenazines in the MWCFC samples with our newer instrument. At the time of our previous analysis, we were using an older HPLC instrument equipped with UV–vis detection but not mass detection, thus we became concerned that we may have mis-identified other compounds as phenazines using this older instrument and decided to revisit our old raw data. In the CHB study, we employed a standard protocol, used by our group and others for routine phenazine analysis in bacteriological cultures, which detects phenazines by their retention times in UV–vis chromatograms (387 nm for pyocyanin and 364 nm for PCA). We used this method to quantify phenazine production from 779 clinical isolates from CF sputum from 47 CHB patients (6). Re-examination of the raw data from these samples confirmed appropriate phenazine assignment. To quantify phenazines in CF sputum, we co-injected pyocyanin or PCA into sputum from which we could not culture *P. aeruginosa* (a major source of phenazines in cystic fibrosis patients) and ran standard curves. In these control co-injection samples, we observed distinct HPLC peaks at 387 nm and 364, as would be expected for pyocyanin or PCA, respectively. Accordingly, we used absorption at these wavelengths to quantify phenazines in all sputum samples. With the benefit of our new awareness that heme B shares retention and absorption characteristics with PCA, we revisited the complete spectral data for each peak from the raw data on our old HPLC, suspecting that we might have unwittingly mis-assigned phenazines to other compounds in sputum. Indeed, analysis of the full UV–vis spectra revealed that the sputum peak we had tracked at 364 nm and assigned to PCA had an absorbance maximum at 398 nm (Figure 2A), suggesting it is unlikely to be PCA despite similar retention times and overlapping absorbance spectra. Instead, this peak’s absorbance spectrum closely matched the spectrum of ferriprotoporphyrin IX identified in our new study (Figure 1D), suggesting we had mis-assigned the heme peak as PCA. Similarly, the sputum peak previously assigned to pyocyanin has an absorbance maximum around 360 nm (Figure 2B) instead of 387 nm, suggesting that pyocyanin is not the major component of this peak. The early-eluting portion of the chromatogram that contains pyocyanin is crowded by multiple overlapping peaks with similar spectra, and so we have been unable to characterize it further.

**Figure 2.**
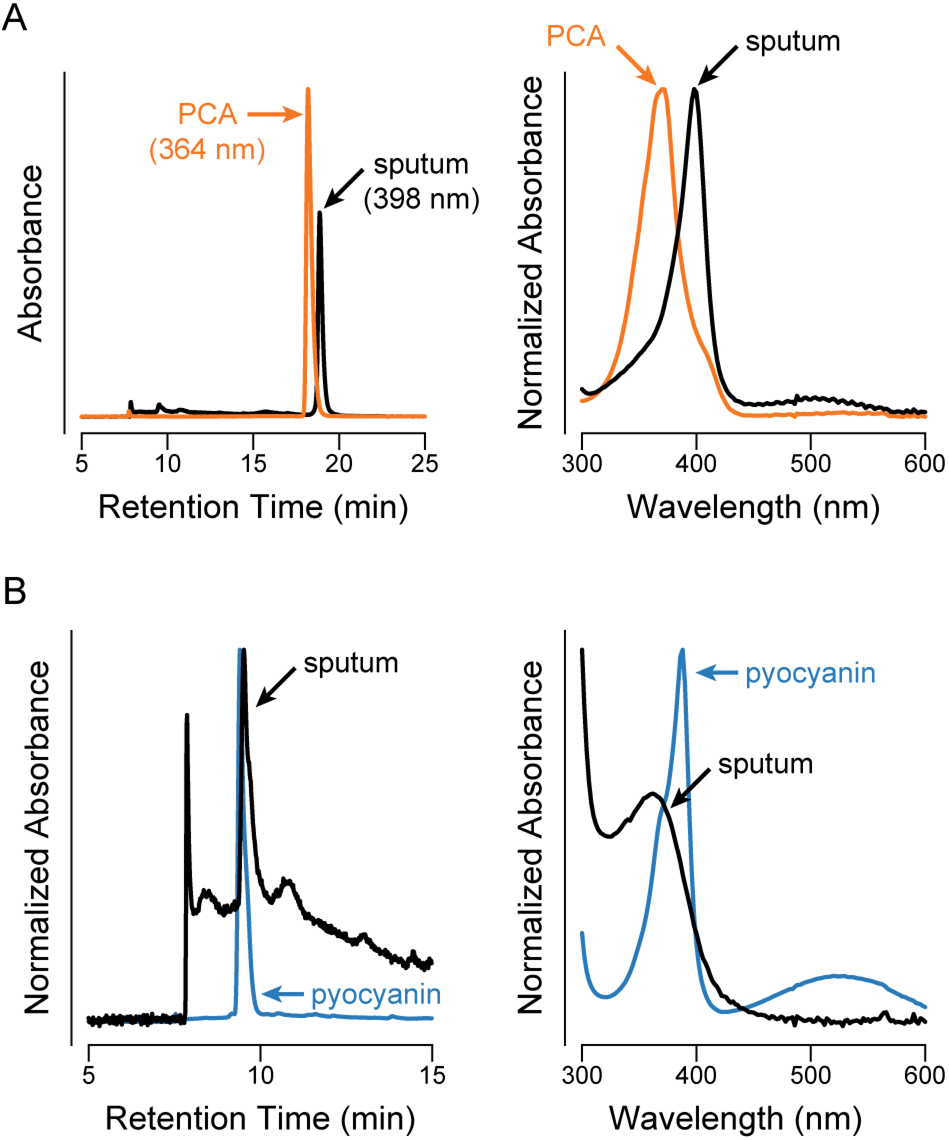
Re-analysis of the HPLC peaks previously assigned as phenazines from the original data collected on an older HPLC. UV–vis chromatograms are shown on the left, and the full UV–vis spectra of the peaks indicated with an arrow are shown on the right. Where indicated on the *y*-axis, the data have been normalized to bring the data into the same scale for clarity. (A) Analysis of the PCA peak, comparing a pure PCA standard (chromatogram at 364 nm, orange) to a sputum sample (chromatogram at 398 nm, black). (B) Analysis of the pyocyanin peak, comparing pyocyanin in *P. aeruginosa*-free sputum (blue) to a *P. aeruginosa*-positive sputum sample (black) (both chromatograms at 387 nm).

Despite our best efforts to perform appropriate controls, including co-injections, the UV–vis spectra reveal that the peaks for pyocyanin and PCA were incorrectly assigned in the original CHB study. This error stemmed from the co-elution of phenazines with sputum compounds that have similar absorbance spectra and the lack of a MS system on the older HPLC instrument. In addition, this instrument was equipped with preparative-scale pumps which led to greater retention time variability between HPLC samples (up to ±1 minute for PCA) compared to analytical instruments, further masking the incorrect assignment. While we cannot rule out the possibility that phenazines were present in our old samples, the distinct UV–vis absorbance spectra indicate that phenazines were not the major component of the assigned peaks, and so our original report overestimated the phenazine concentration in CF sputum samples.

### Heme concentrations correlate negatively with lung function

As the re-analysis suggests, the interpretation of our original HPLC data was confounded by coeluting compounds that are present in sputum. The similarity between the peak we previously analyzed as PCA and what we have now identified as ferriprotoporphyrin IX (Figure 1D) suggests that, instead of a correlation between phenazines and FEV_1_ %, the original study measured a correlation between heme and FEV_1_%. To evaluate this possibility, we quantified the concentration of ferriprotoporphyrin IX in our MWCFC samples from our new dataset. A plot of heme concentration versus FEV_1_% reveals a statistically significant (Spearman’s ρ = −0.47, *p* < 0.001) correlation between sputum heme levels and FEV_1_% (Figure 3). Three of the samples contained substantially more heme than the others, and this correlation held even when the outliers with >10 µM heme were excluded from the analysis (ρ = −0.41, *p* < 0.001) because the Spearman rank correlation is independent of the magnitude of the variables. Although the concentration of ferriprotoporphyrin IX in the original samples is unknown, it is likely that the original PCA concentration data instead represents heme concentration with an unknown conversion factor, and so both studies have independently confirmed a correlation between the heme signal in HPLC analysis and lung function in cystic fibrosis patients.

**Figure 3.**
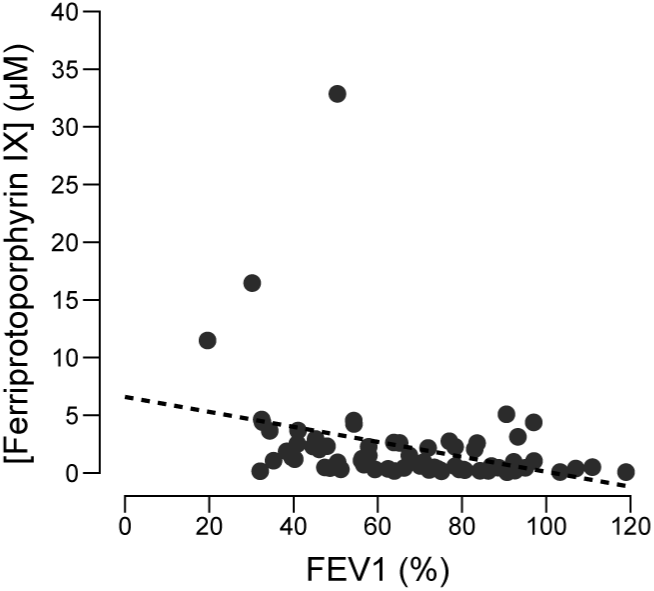
Correlation between ferriprotoporphyrin IX concentration and disease status as measured by percent predicted FEV_1_ (Spearman’s ρ = −0.47, *p* = 3.6 × 10^−5^). Each data point represents a single sputum sample from a unique patient. The dashed line illustrates a linear regression through the data.

## Discussion

Several independent analyses have detected phenazines in sputum from cystic fibrosis patients (4, 5). Our new data collection effort and the correction reported here to our previous results (6) was partially motivated by a more recent report of detectable phenazine concentrations in only 3 of 27 patients (4). Interestingly, extensive isolation of several hundred clinical *P. aeruginosa* strains in our lab has verified that phenazine production is a nearly universal capability of *P. aeruginosa* strains that inhabit cystic fibrosis patients, a result that is still valid from our original study (6). All phenazine analyses in the original paper that were performed in bacteriological growth medium were accurate. We therefore find it interesting that phenazines and other *P. aeruginosa* metabolites are detected only sporadically in sputum samples, often from patients that are culture-negative for *P. aeruginosa* (4). Together these findings are suggestive of significant macro- and micro-scale heterogeneity within cystic fibrosis lungs. Consistent with regional heterogeneity, *P. aeruginosa* strains within different regions are known to undergo independent evolution with minimal cross-colonization (11). It is difficult to verify that independent samples originate from the same region of the lung, and so heterogeneity between sputum samples may account for differences between metabolite and microbial analyses within the same patient. Furthermore, sputum samples are extremely viscous and poorly mixed, leading to high spatial heterogeneity within individual samples (12). Sputum viscosity originates from extracellular polymers including DNA (13), and because pyocyanin and other phenazines are known DNA intercalators (14), it is likely that cells within the sputum matrix encounter a micro-scale phenazine concentration that is considerably higher than that determined by bulk measurements. We anticipate that new methods to image metabolites at large and small spatial scales will provide insights into the cystic fibrosis lung environment and clarify the role of pathogen-derived metabolites in disease progression. Although it remains possible that phenazine concentration negatively correlates with lung function, our original study overestimated the total phenazine concentration and the true concentrations were likely below our limit of detection.

The form of heme we measured in this study, ferriprotoporphyrin IX or heme B, is the most abundant heme found in humans and the oxygen-carrying component of hemoglobin in red blood cells. Given that the majority of components in lung sputum are host-derived (4), the heme likely originated from blood rather than from microbial sources. Consistent with this assumption, hemoptysis, or coughing up blood, is observed in up to 60% of cystic fibrosis patients (15–17). While minor blood streaking in sputum does not usually prompt treatment, massive hemoptysis is a marker of pulmonary exacerbations and correlates with impaired lung function and mortality in adults (15, 16), and so the correlation between heme and lung function decline may identify heme as a potentially useful biomarker of clinical disease in CF.

Previously we also reported a correlation between sputum iron and lung function decline (18). This correlation arose from the abundance of Fe(II) which predominated, rather than from Fe(III) (18). We also reported a correlation between Fe(II) and what we believed was PCA (18), which we now recognize was heme. As heme is a major carrier of iron, this correlation suggests blood-derived heme may be a significant source of sputum iron. In the context of lung infections, iron is an important nutritional regulator of pathogenicity and biofilm formation in bacteria, including *P. aeruginosa* (19), and pathogenic bacteria can extract iron from heme (9). Given that iron chelation is now being explored as a potential therapy for lung infections (20), there is a need to better understand the sources of sputum iron and its influx rate, of which our results suggest host-derived heme could be a significant contributing factor.

The prognosis for cystic fibrosis patients has improved remarkably in recent decades, owing not only to advances in treatments but also in diagnostics and preventative care (21). While the role of pathogen-derived metabolites in disease state is still uncertain, host-derived factors are clearly abundant (4) and may provide new markers to guide treatment. The prevalence of heme in every sputum sample we measured, and the simplicity of its detection by HPLC owing to its characteristic 398 nm absorbance peak, suggest it can be a useful diagnostic component for monitoring cystic fibrosis disease progression. Additional research may uncover a causative link between sputum heme and lung disease through the interplay between iron and pathogenic microbes.

## Materials and Methods

### HPLC method from the original study

The original study used a Beckman Gold HPLC equipped with preparative-scale pumps on a model 126P solvent module. The separation used a Waters Symmetry C18 column with dimensions 4.6 mm × 250 mm and a particle size of 5 µm. The injection volume was 100 µl, supplied via a model 508 autosampler, and the flow rate was 1.0 mL/min. Solvent A was water with 0.1% trifluoroacetic acid (TFA) and Solvent B was acetonitrile with 0.1% TFA. Compounds were eluted using a linear gradient of 15% B from 0 to 2 minutes, 15% to 83% B from 2 to 22 minutes, and 0% B from 22 to 30 minutes. UV–vis data were collected from 200 to 600 nm using a model 168 photodiode array detector.

### Sample preparation

Sputum samples were obtained after informed consent from patients with confirmed CF aged 6 and older able to expectorate sputum during clinical stability. The study protocol was reviewed and approved at each of the 9 participating centers in the Mountain West CF Consortium (MWCFC). Patients were randomly selected from the MWCFC Centers in order to enroll a representative sample of all US patients with CF. Samples were processed within 4 hours of expectoration by dilution 1:1 with Hanks buffered saline solution (Sigma, St Louis, MO), vigorous vortex mixing for 1 minute followed by centrifugation at 2,800 g and 4 °C for 20 minutes to obtain lipid and aqueous fractions and pellets. For this study, we used aqueous fractions that were mixed 1:1 with protease inhibitor cocktail (Sigma). Fractions were frozen and stored at −80 °C until use. The samples were thawed at room temperature and centrifuged at 13,000 *g* for 15 minutes. The supernatant was transferred to a Waters Total Recovery Vial and injected directly into the HPLC without further processing. Hemin was purchased from Sigma-Aldrich and was dissolved to a stock concentration of 10 mM using 100 mM aqueous ammonia. A standard curve from 0.01 to 100 µM was created by diluting the stock solution into water.

### Spirometry

Clinical data including spirometry data were collected at the same time as sputum samples. The forced expiratory volume in 1 second (FEV_1_) was measured according to American Thoracic Society standards (22). FEV_1_ was normalized to percent predicted FEV_1_ (FEV_1_%) for age, sex, height, race and ethnicity using NHANES III equations (23) as was done clinically at all participating centers at the time of study enrollment.

### High resolution LC-MS

HPLC separations for high resolution mass spectrometry were performed on a Waters Acquity I-Class UPLC connected to a Waters Acquity PDA detector and a Waters Xevo G2-XS quadrupole time-of-flight mass spectrometer. The sample chamber was maintained at 4 °C. The separation used a Waters XBridge BEH C18 XP column with dimensions 3.0 × 100 mm and a particle size of 2.5 µm. The column was equipped with a VanGuard guard column comprising the same resin. The injection volume was 5 µl and the flow rate was 0.5 mL/min. The column was maintained at 40 °C. Solvent A was water with 0.1% formic acid and solvent B was acetonitrile with 0.1% formic acid. Compounds were eluted using a linear gradient of 0% to 90% from 0 to 24 minutes, 90% B from 24 to 25 minutes, and 0% B from 25 to 35 minutes. Mass scans from 150 to 1000 Da were collected in separate chromatographic runs for positive and negative ionization modes with a scan time of 0.3 s. For both modes, the mass spectrometry parameters were: capillary voltage, 1.5 kV; sampling cone, 30 V; source offset, 80 V; source, 120 °C; desolvation, 550 °C; cone gas, 50 L/h; desolvation gas, 800 L/h. The analyser mode was set to resolution and the dynamic range was set to normal. In the negative ion mode, fragmentation data were collected in a second data channel using a collision-ramp energy of 10 to 14 eV. A solution of 20 pg/μL leucine enkephalin was used as the LockSpray solution with a flow rate of 20 μl/min. The instrument was controlled using the software MassLynx.

### Routine LC-MS for ferriprotoporphyrin IX quantification

For routine analysis, HPLC separations were performed on a Waters Alliance HPLC connected to a Waters 2998 PDA detector and a Waters Acquity QDa single quadrupole mass spectrometer. The sample chamber was maintained at 10 °C. Separations were performed the same as for the high-resolution method, except that the gradient included a constant 2% methanol supplied via solvent C, which was found to greatly improve the reproducibility of pyocyanin measurements. Mass scans were collected simultaneously from 100 to 900 Da in the positive mode and 100 to 600 Da in the negative mode. In addition, a selected ion recording channel was collected in the positive mode for 211.09, 224.08, and 225.07 Da, the masses expected for pyocyanin, phenazine-1-carboxamide, phenazine-1-carboxylic acid, respectively. The probe temperature was 600 °C and the capillary voltage was 0.8 kV. UV–vis data were collected from 200 to 800 nm at 20 Hz with a resolution of 1.2 nm. The instrument was controlled using the software Empower. Peaks representing ferriprotoporphyrin IX were integrated using the Apex Track algorithm within Empower and compared against a linear calibration curve of samples derived from hemin. The measured concentrations were multiplied four-fold to account for the dilution that occurred during sample processing.

### Data analysis and statistics

To ensure uniformity, data from the three instruments were exported and compared together within a series of Python scripts. Plots were generated using the Matplotlib package and labeled with Adobe Illustrator. The Spearman correlation coefficient was calculated using the spearmanr function in the SciPy package.

## Acknowledgments

The MWCFC Sputum Biomarker investigators included Frederick R Adler, Fadi Asfour, Barbara A Chatfield, Jessica A Francis, John R Hoidal, Judy L Jensen, Yanping Li, Theodore G Liou, Kristyn A Packer, Jane Vroom (University of Utah); Natalia Argel, Peggy Radford (Phoenix Children’s Hospital); Perry S Brown, Dixie Durham (St. Luke’s Cystic Fibrosis Center of Idaho); Cori L Daines, Osmara Molina (University of Arizona); Barbara Glover, Craig Nakamura, Ryan Yoshikawa (Cystic Fibrosis Center, Las Vegas); Theresa Heynekamp, Abby J Redway (University of New Mexico); Ruth Keogh (London School of Hygiene and Tropical Medicine); Carol M Kopecky, Scott D Sagel (Children's Hospital Colorado, University of Colorado School of Medicine); Noah Lechtzin (Johns Hopkins University School of Medicine); Jerimiah Lysinger, Shawna Sprandel (Montana Cystic Fibrosis Center, Billings Clinic); Katie R Poch, Jennifer L Taylor-Cousar (National Jewish Health); Alexandra L Quittner (Miami Children's Research Institute, Nicklaus Children's Hospital); John P Clancy (University of Cincinnati); J Stuart Elborn (Queen’s University, Belfast and Royal Brompton Hospital, London); Kenneth N Olivier (Laboratory of Chronic Airway Infection, Pulmonary Branch, National Heart Lung and Blood Institute, National Institutes of Health). We thank Dr. Nathan Dalleska at the Environmental Analysis Center (Caltech) for analytical support.

